# Hysteresis and noise floor in gene expression optimised for persistence against lethal events

**DOI:** 10.1101/2024.07.19.604229

**Authors:** Pavol Bokes, Abhyudai Singh

## Abstract

Bacterial cell persistence, crucial for survival under adverse conditions like antibiotic exposure, is intrinsically linked to stochastic fluctuations in gene expression. Certain genes, while inhibiting growth under normal circumstances, confer tolerance to antibiotics at elevated expression levels. The occurrence of antibiotic events lead to instantaneous cellular responses with varied survival probabilities correlated with gene expression levels. Notably, cells with lower protein concentrations face higher mortality rates. This study aims to elucidate an optimal strategy for protein expression conducive to cellular survival. Through comprehensive mathematical analysis, we determine the optimal burst size and frequency that maximise cell proliferation. Furthermore, we explore how the optimal expression distribution changes as the cost of protein expression to growth escalates. Our model reveals a hysteresis phenomenon, characterised by discontinuous transitions between deterministic and stochastic optima. Intriguingly, stochastic optima possess a noise floor, representing the minimal level of fluctuations essential for optimal cellular resilience.

## 1 Introduction

Noisy gene expression constitutes a significant source of phenotypic variability within populations of genetically identical cells [1, 2]. This heterogeneity arises from the inherent discreteness of transcription and/or translation reactions [3, 4]. Mathematical modeling of these processes elucidates the relationship between kinetic gene expression parameters and the resulting population distribution of protein concentration [5–7]. Notably, even highly abundant proteins display a basal level of noise, commonly known as the noise floor [8, 9].

Stochastic gene expression is effectively modeled by a framework rooted in the concept of bursty protein synthesis [10, 11]. Experimental evidence has documented gene expression bursts at both transcriptional and translational levels [12, 13]. This bursty model accurately predicts a gamma distribution of protein concentration within cellular populations [14, 15]. Notably, the shape and scale parameters of this distribution directly correspond to the mean frequency and size of bursts, providing valuable insights into the gene expression dynamics [16, 17].

Phenotypic variability leads to the persistence of clonal cells following treatment [18, 19]. This phenomenon entails a fractional killing of the cell population, wherein a portion of sensitive cells succumbs while a persistent fraction survives [20]. Notably, persistent cells are not characterized as resistant mutants and may retain sensitivity to subsequent treatments [21, 22]. Bacterial persistence has been correlated with heightened toxin expression and diminished cell growth rates [23–25] (see Figure 1). Notably, protein-dependent suppression of cell growth limits protein dilution and constitutes a positive feedback loop on protein expression [26, 27]. Trade-off between growth and persistence is also observed in cancer cells [28, 29].

**Figure 1:**
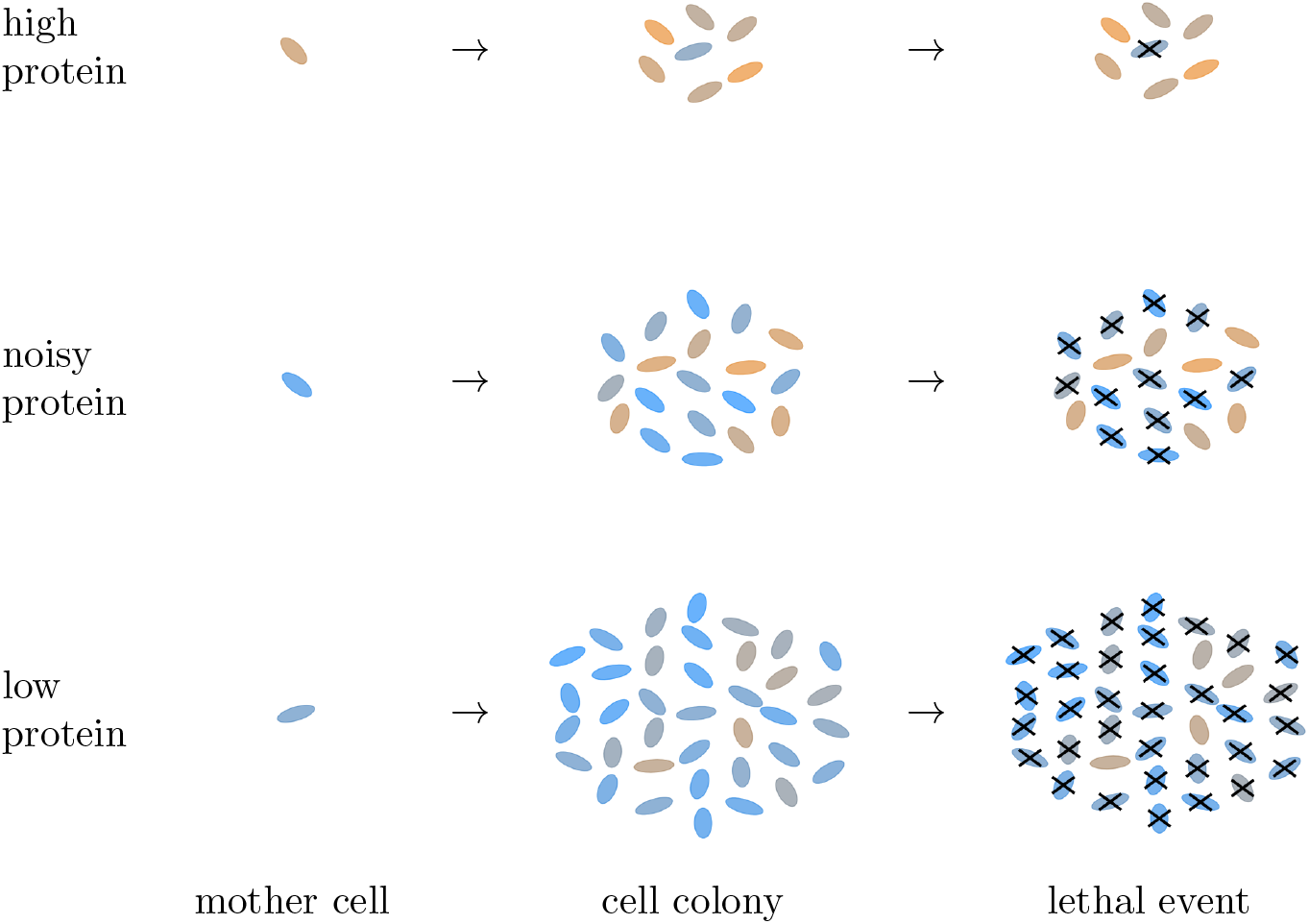
High protein expression imposes a cost to cell growth but provides protection at lethal events. Low protein expression enables growth but is sensitive to lethal events. The aim is to find optimal protein expression parameters (burst size and frequency) that maximise the expected colony size post lethal event.

Evolutionary pressures drive the selection of gene expression strategies that maximize species growth. Neglecting population heterogeneity may result in the adoption of suboptimal strategies [30, 31]. Achieving optimal gene expression involves assessing the trade-off between the cost of protein production and the benefits of protein-dependent growth [32] such as improved precision at high protein levels [33]. Other investigations have focused on maximizing mutual information between a transcription factor and the expression level of its target gene [34, 35]. Additionally, synthetic gene regulatory circuits have been proposed to sense and respond to environmental changes, adjusting their activity for optimal performance [36].

**Table 1:** Model parameters and variables. Without loss of generality, unit time is chosen so that *r*(0) = 1; unit protein concentration is set so that *s*(1) = (*s*_0_ + *s*_*∞*_)*/*2.

| Notation | Interpretation |
| --- | --- |
| $y$ | random process representing intercellular protein concentration |
| $n(y, t)$ | number of cells with protein concentration $y$ at time $t$ |
| $r(y)$ | cell growth rate as function of protein concentration $y$ |
| $a$ | burst frequency — the rate of occurrence of protein bursts |
| $b$ | the mean size of exponentially distributed protein bursts |
| $x = ab$ | steady state mean protein concentration |
| $k$ | cost to cell growth rate per unit protein concentration |
| $T$ | mean waiting time until a lethal event |
| $\mu = kT$ | protein burden on growth at the time of a lethal event |
| $s(y)$ | survival probability as function of protein concentration |
| $s_0$ | probability that a cell without protein survives a lethal event |
| $s_\infty$ | survival probability of a cell with an infinite amount of protein |
| $h$ | Hill coefficient for the survival probability function |
| $f(x, b)$ | cost of protein expressed at mean $x$ in bursts of mean size $b$ |

This paper centers on elucidating the role of a protein whose expression confers antibiotic tolerance while hampering cell growth. Section 2 presents the formulation of the bursty gene expression model alongside the associated cost function for the protein. In Section 3, we identify the critical points of this cost function and classify their type. Subsequently, in Section 4, we conduct a bifurcation analysis of these critical points with respect to a parameter measuring the burden that the protein places on cell growth. Finally, in Section 5, we offer a comprehensive summary and interpretation of our findings.

## 2 Model formulation

We consider a model for stochastic gene expression in a growing population of cells. We assume that a protein is produced in each cell in bursts as per a Poisson process with burst frequency *a*. In each burst, a random amount of protein is produced, which we assume to be exponentially distributed with mean size *b*. Cells proliferate with rate *r* = *r*(*y*) which depends on the cellular concentration *y* of the protein. Cell growth continually dilutes the protein concentration as per the non-linear decay equation *dy/dt* = −*yr*(*y*). The number *n*(*y, t*) of cells with protein concentration *y* at time *t* satisfies an integro-partial-differential equation

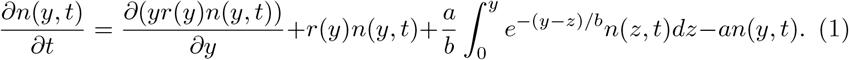

This population balance equation holds in a suitable time interval and for *y* > 0; the point *y* = 0 is a natural boundary. The four terms on the right-hand side of (1) quantify the change in population composition due to protein dilution, cell growth, burst influx, and burst efflux, respectively.

A single-cell version of the equation without the growth term was introduced in [14] and subsequently studied by other authors (e.g. [37, 38]) including us [39, 40]. We and co-authors extended the model by protein-dependent growth and analysed differences between the single-cell and population models in [41, 42]. The difficulty introduced by the growth term can be recognised by integrating (1) over all admissible protein concentration values *y* ∈ (0,∞). Doing so shows that the total cell count

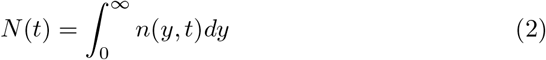

satisfies

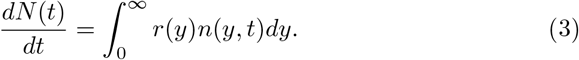

If the growth rate *r*(*y*) is independent of the protein level *y*, equation (3) is closed and immediately implies exponential growth. Otherwise, (3) is not closed and subtler methods are required to calculate the principal eigenvalue [42].

Here we circumvent this difficulty by assuming that the effect of the protein is relatively small:

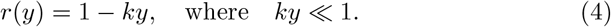

Without loss of generality, the growth rate is set to one in the absence of the protein.

After an initial transient, solutions to (1) are approximately equal to

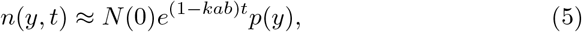

where

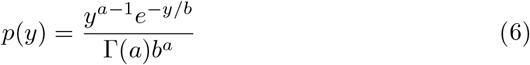

is the normalised protein distribution (the gamma distribution). We note that the gamma distribution is independent on the burden *k* imposed by the protein. The effect of the burden is in slowing down the protein growth by a factor proportional to the steady state protein mean *ab*. These results are consistent with the exact analysis in [42]. We emphasise that this simplicity is due to the assumption of a small burden in (4). Without this assumption, the differentials in growth rates at different protein concentrations will tilt the protein probability distribution toward faster growing cells [42].

Cells that express the protein proliferate slower but have a better chance of survival of stress events (treatment). The probability of surviving treatment in the presence of *y* protein is modelled by a sigmoid fuction

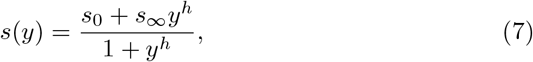

with 0 ≤ *s*_0_ < *s*_*∞*_ and *h* > 1 assumed. The protein concentration at which the survival probability is equal to the average value (*s*_0_ + *s*_*∞*_)*/*2 is thereby fixed to *y* = 1 without loss of generality. The proportion of cells surviving treatment is given by the integral

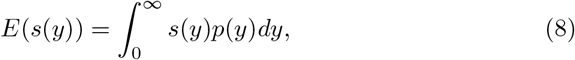

where the protein probability density function *p*(*y*) depends on the gene expression parameters *a* and *b* as specified in (6).

We assume that treatment events occur as per a Poisson process with mean interarrival times *T*. Over a typical period from one treatment to another, the population grows by the factor of

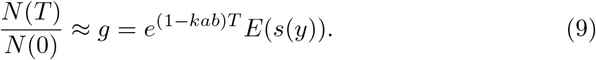

The fittest cells maximise *g* with respect to the parameters of the gene expression model. We apply on the dependent variable a strictly decreasing transformation *f* = *T*−ln *g*. Our goal is therefore to minimise with respect to the protein mean *x* = *ab* and the mean burst size *b* the *cost function*

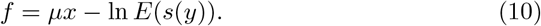

The product *µ* = *kT* is henceforth referred to as *protein burden rate*. Dividing *s*(*y*) by *s*_*∞*_ is equivalent to adding a costant term ln *s*_*∞*_ to the cost function; therefore, *s*_*∞*_ = 1 can be assumed without loss of generality.

We parametrise the cost function by the protein mean *x* = *ab* > 0 and burst size *b* > 0. On the boundary of the parameter domain, the following asymptotic behaviour is identified (cf. Figure 2):

**Figure 2:**
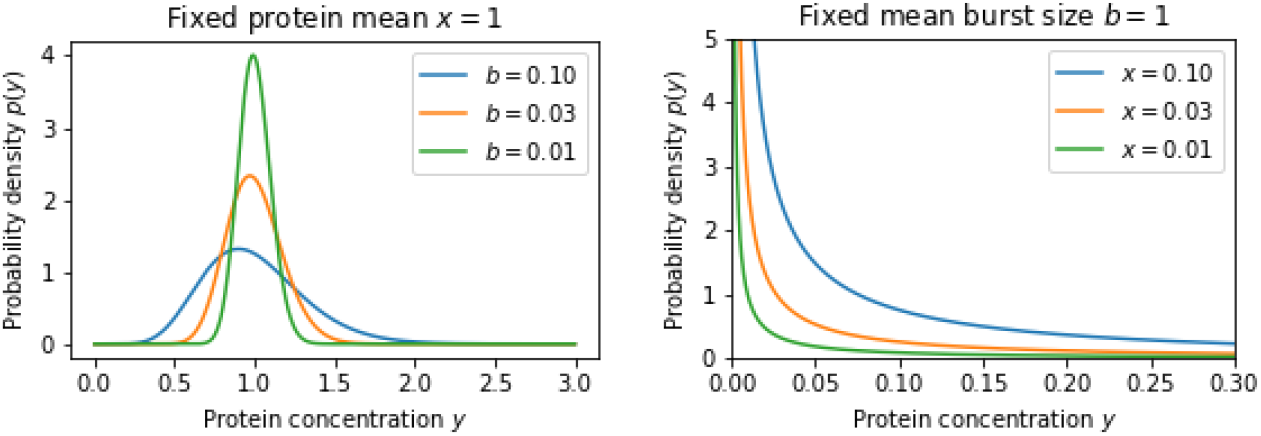
Protein distribution (6) for varied values of protein mean *x* and expected burst size *b*

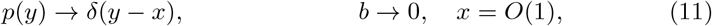

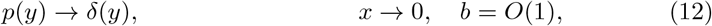

where *δ*(*y*) denotes the Dirac delta function. Regime (11) corresponds to a frequently expressed (*a* = *x/b*→∞) protein which deviates little from its mean value *x*. Regime (12) corresponds to a protein expressed rarely (*a* = *x/b*→0) in random bursts of mean size *b*. Both regimes will play a role in the optimisation problem.

The above asymptotics allow to extend the distribution (6) to the closure *x* ≥ 0 and *b* ≥ 0 of the parameter domain. For *x* ≥ 0 and *b* = 0, the limit in (11) gives a deterministic protein expression state; increasing *b* from zero to a small positive value while keeping *x* unchanged makes a small, nearly Gaussian, stochastic perturbation to the deterministic expression. For *x* = 0 and *b* ≥ 0, the limit (12) gives the non-expression state of the protein; increasing *x* from zero to a small positive value while keeping *b* unchanged makes a highly skewed and non-Gaussian perturbation of the non-expression state. These types of perturbations will be useful for characterising the character of the deterministic/non-expression states in the optimisation problem.

## 3 Critical points of the cost function

We want to minimise the cost function *f* = *f* (*x, b*) as function of protein mean *x* and burst size *b* within the positive quadrant *x* ≥ 0, *b* ≥ 0. More generally, we are also interested in the critical points of *f* (*x, b*), which in addition to local minimisers also include maximisers and saddle points. Leaving aside the non-expression state, critical points can be deterministic (*b* = 0) or stochastic (*b* > 0).

### 3.1 Deterministic critical points

Inserting (11) into (8) gives *E*(*s*(*y*)) = *s*(*x*) for *b* = 0. The cost of expressing the protein deterministically at level *x* is thus equal to

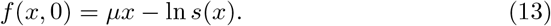

By a deterministic critical point we understand a point *x* > 0 and *b* = 0 at which the cost is stationary:

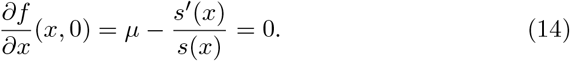

The character of the critical point is determined by the partial derivatives

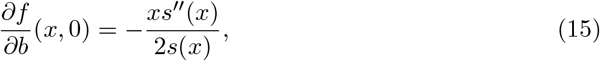

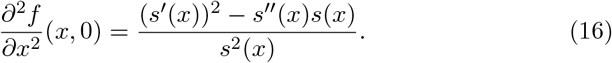

Equation (16) is obtained by differentiating (14) with respect to *x*. If (16) is positive (negative), then perturbing the critical point by a small deterministic value increases (decreases) the cost. Equation (15) follows from a linear noise approximation (Appendix A). If (15) is positive (negative), then adding a small amount of stochastic noise increases (decreases) the cost. Clearly, the critical point is a point of (i) minimum if 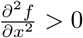 and 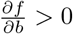 (ii) maximum if 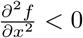 and 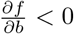 (iii) saddle if 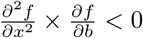.

### 3.2 Stochastic critical points

By a stochastic critical point we understand a point *x* > 0 and *b* > 0 at which

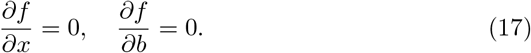

This is further distinguished depending on the Hessian matrix

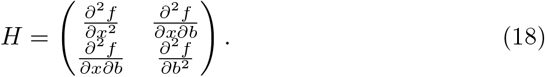

A stochastic critical point is a point of (i) minimum if *H* is positive definite; (ii) maximum if *H* is negative definite; (iii) saddle if *H* is indefinite.

## 4 Bifurcation analysis

We are interested in the problem of parameter continuation, i.e. determining how the coordinates and the character of critical points evolve as a bifurcation parameter changes. We use the protein burden rate *µ* as the bifurcation parameter and denote by *µ*_1_ < *µ*_2_ < … its bifurcation values. We also denote by *x*_1_, *x*_2_, … and *b*_1_, *b*_2_ … the coordinates of the bifurcation points. The relatively simpler situation is treated first which occurs when survival is impossible without any protein (*s*_0_ = 0); the effects of introducing a basal chance of survival (*s*_0_ > 0) are studied thereafter. The latter case is referred to as “leaky” and the former as “tight”.

### 4.1 Tight case (*s*_0_ = 0)

By (14) and (7), deterministic critical points satisfy

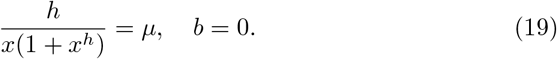

The left-hand side of (19) is a decreasing function of *x*, approaching infinity as *x*→0 and diminishing to zero as *x*→∞ Therefore, there exists a unique deterministic critical point *x* = *x*(*µ*), *b* = 0 for any given *µ* > 0. This is shown in the bifurcation diagrams of Figure 3 in red colour.

**Figure 3:**
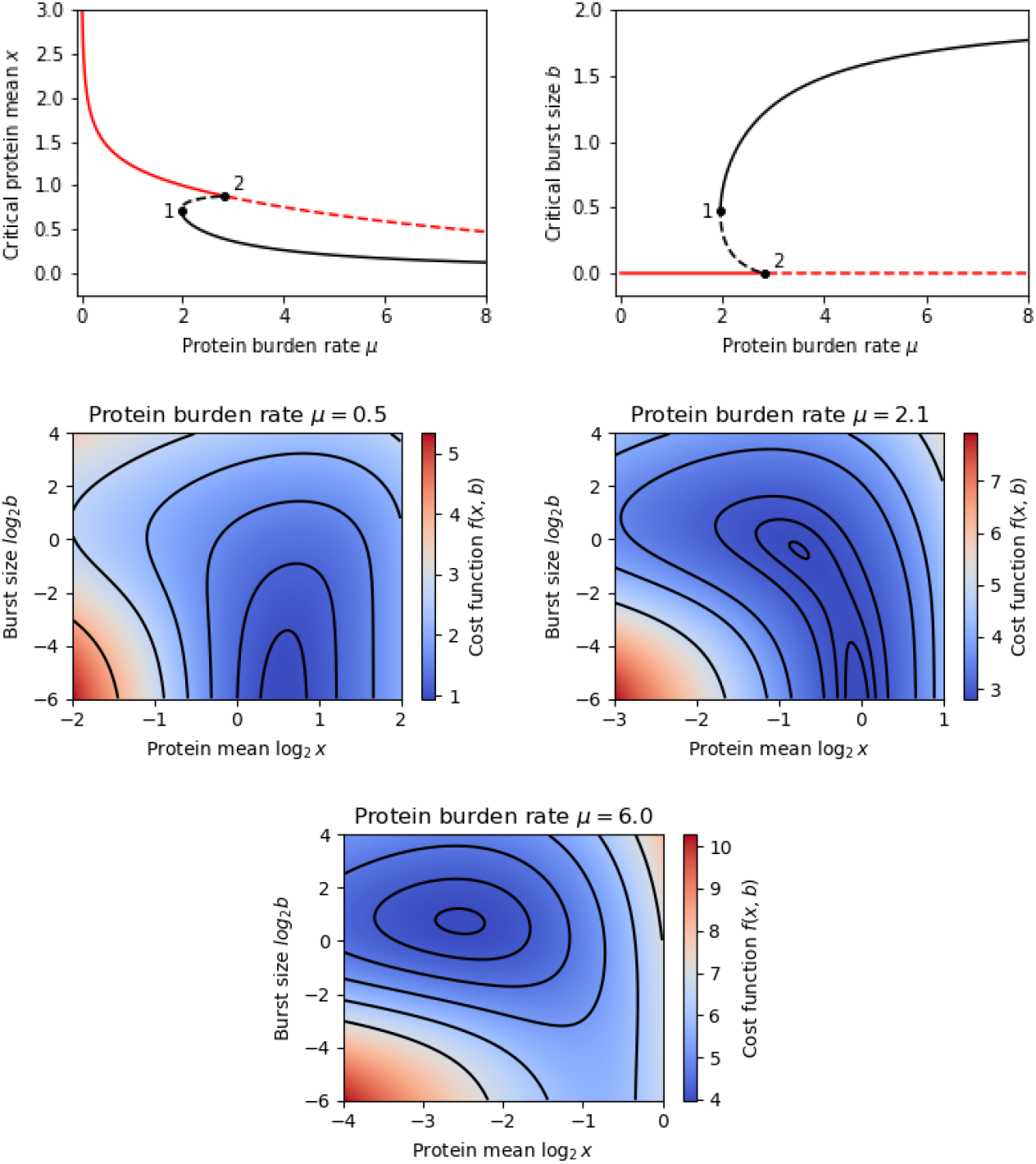
*Top panels:* The *µ*-dependence of the *x* and *b* coordinates of the cost function minimisers (full lines) and saddle-points (dashed lines) in the tight case (*s*_0_ = 0). Markers indicate the bifurcation points. *Central and bottom panels:* The cost function *f* (*x, b*) over a range of (logarithmically scaled) protein means *x* and burst sizes *b* for selected values of the protein burden rate *µ. Other parameter values: s*_*∞*_ = 1, *h* = 4.

The character of the deterministic critical point is determined by the signs of (15) and (16). Monotonicity of (19) implies that (16) is positive. Depending on the sign of (15), the deterministic critical point can be a minimum or a saddle point. Equation (15) simplifies to

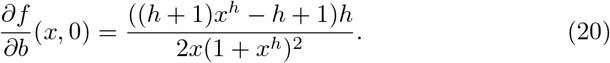

Equating the numerator of (20) to zero and inserting into (19) give the explicit bifurcation values (cf. Figure 3)

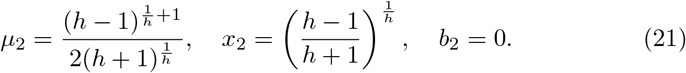

The critical point is a minimum if *µ* < *µ*_2_ (i.e. *x* > *x*_2_) and a saddle point if *µ* > *µ*_2_ (i.e. *x* < *x*_2_).

In addition to the deterministic critical point, we can also have up to two stochastic critical points for any value of *µ*. The *µ*-dependence is determined numerically using the method of Appendix B. The overall bifurcation picture as follows (cf. Figure 3): for *µ* < *µ*_1_ = 1.97 the only critical point is the deterministic minimum. A typical heat map of *f* (*x, b*) is depicted in central left panel. At *µ* = *µ*_1_, a saddle-node bifurcation occurs at (*x*_1_, *b*_1_) = (0.71, 0.48). For *µ*_1_ < *µ* < *µ*_2_ = 2.84, a stochastic minimum and a stochastic saddle point exist in addition to the deterministic minimum. A typical heat map of *f* (*x, b*) is depicted in central right panel. At *µ* = *µ*_2_, the stochastic saddle coalesces with the deterministic minimum in a transcritical bifurcation. For *µ* > *µ*_2_, there is a deterministic saddle and a stochastic minimum. A typical heat map of *f* (*x, b*) is depicted in the bottom panel.

### 4.2 Leaky case (*s*_0_ > 0)

If *s*_0_ > 0, the function *s*′(*x*)*/s*(*x*) is non-monotonous, and equation (14) for deterministic critical points has two solutions if *µ* < *µ*_3_, one solution if *µ* = *µ*_3_, and no solution if *µ* > *µ*_3_, where *µ*_3_ is a point of a saddle-node bifurcation. The *µ*-dependence of the deterministic critical points is shown in red colour in Figure 4.

**Figure 4:**
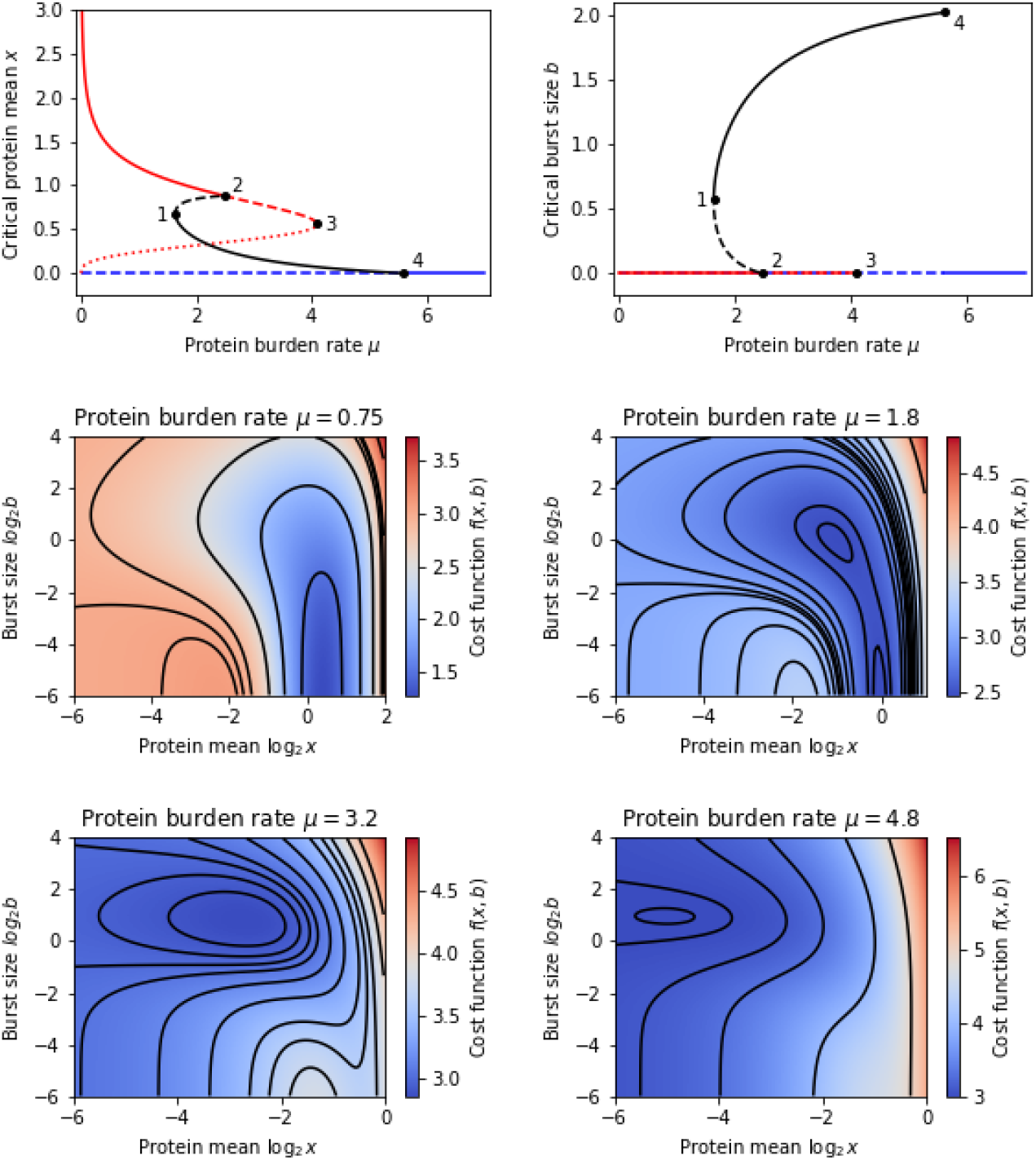
*Top panels:* The *µ*-dependence of the *x* and *b* coordinates of the cost function minimisers (full lines), saddle-points (dashed lines), and maximisers (dotted line) in the leaky case (*s*_0_ = 0.05). Markers indicate the bifurcation points. *Central and bottom panels:* The cost function *f* (*x, b*) over a range of (logarithmically scaled) protein means *x* and burst sizes *b* for selected values of the protein burden rate *µ. Other parameter values: s*_*∞*_ = 1, *h* = 4.

If *s*_0_ > 0, we also need to consider the trivial critical point with *x* = 0 (no protein expression) which has a finite cost

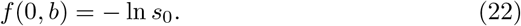

The character of the trivial critical point is determined by the sign of

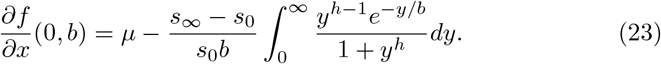

Formula (23) follows from an asymptotic evaluation of the cost function (Appendix A). In Appendix A, we also show that there exists a bifurcation value *µ*_4_ such that (i) if *µ* < *µ*_4_, then *∂f* (0, *b*)*/∂x* changes sign in the interval 0 ≤ *b* < ∞; (ii) if *µ* > *µ*_4_, then *∂f* (0, *b*)*/∂x* > 0 for all *b* ≥ 0. Therefore, we classify the trivial critical point as a saddle point if *µ* < *µ*_4_ and a minimum if *µ* > *µ*_4_.

The overall bifurcation behaviour is as follows (cf. Figure 4): for *µ* < *µ*_1_ = 1.63, there are two deterministic critical points (a maximum and a minimum) and the trivial critical point, which is a saddle point. A typical heat map of *f* (*x, b*) is depicted in the central left panel. At *µ* = *µ*_1_ = 1.63, a saddle–node bifurcation occurs at (*x*_1_, *b*_1_) = (0.67, 0.57). It is the same kind of bifurcation that is observed in the tight case, but the bifurcation values are perturbed due to the leakiness. For *µ*_1_ < *µ* < *µ*_2_ = 2.48, a stochastic minimum and a stochastic saddle point exist in addition to the aforementioned deterministic critical points. A typical heat map of *f* (*x, b*) is depicted in the central right panel. At *µ* = *µ*_2_, the stochastic saddle point and the deterministic minimum collapse into a deterministic saddle point. Again, the same kind of transcritical bifurcation occurred in the tight case, but at a slightly different value of the bifurcation parameter. For *µ*_2_ < *µ* < *µ*_3_ = 4.1, there is a stochastic minimum, a deterministic maximum, a deterministic saddle point, and the trivial saddle point. A typical heat map of *f* (*x, b*) is depicted in the bottom left panel. At *µ* = *µ*_3_, the deterministic saddle point and maximum coalesce in a saddle-node bifurcation. This bifurcation does not have an analogue in the tight case. For *µ*_3_ < *µ* < *µ*_4_ = 5.61, we have the trivial saddle point and a stochastic minimum. A typical heat map of *f* (*x, b*) is depicted in the bottom right panel. At *µ* = *µ*_4_, the stochastic minimum coalesces with the trivial saddle point in a transcritical bifurcation. This bifurcation also does not have an analogue in the tight case. For *µ* > *µ*_4_, the only critical point is the trivial one, which is a minimum.

## 5 Conclusion

We considered a stochastically expressed protein which confers tolerance to antibiotic treatment but places a burden on the cell machinery and inhibits cell growth. The population growth observed in the presence of such a protein is quantified by a population balance equation. The underlying assumption behind the use of the equation is that the population density is continuous; it does not account for discrete population effects such as extinction and fixation [43]. In addition to the continuity assumption, we assumed that (i) the cost of protein to growth is relatively small and (ii) the waiting times till treatment are relatively large. Owing to these assumptions, the population distribution can be assumed to have equilibrised at a gamma distribution with shape and scale directly linked to the protein burst frequency and the burst size. Without these simplifying assumptions, we expect that the analysis would have to rely on a numerical solution of the integro-partial-differential population balance equation.

Subject to the aforementioned simplifications, the overall effect of the protein on the cell growth is quantified by a tractable cost function *f*. The cost function depends on the growth parameters (protein burden rate *µ*; survival probabilities *s*_0_ and *s*_*∞*_; the Hill constant *h*) and the gene expression parameters (protein mean *x*; mean burst size *b*). Gene expression can be stochastic (*x* > 0 and *b* > 0), deterministic (*x* > 0, *b* = 0), or turned off (*x* = 0, *b* 0).

The cost is minimised as function *f* = *f* (*x, b*) of the gene expression parameters for a given set of growth parameters. Depending on how cells without protein respond to treatment, we differentiated between the tight case (*s*_0_ = 0) and the leaky case (*s*_0_ > 0). Optimal gene expression can be stochastic or deterministic; in the leaky case it can be optimal not to express any protein.

The main result of our analysis is the description of how optimal gene expression parameters change in response to changing protein burden rate *µ*. This is visualised in bifurcation diagrams in Figure 3 (tight case) and Figure 4 (leaky case). In addition to minima, the bifurcation diagrams also show the *µ*-dependence of other types of critical points of the cost function, namely saddle points and maxima, and highlight bifurcation points *µ*_1_ < *µ*_2_ < *µ*_3_ < *µ*_4_, at which the critical points coalesce and change type.

Increasing protein burden rate *µ* leads to a decrease in the mean value *x* of the optimal protein expression. In the tight case, there is a single deterministic optimum if *µ* < *µ*_1_, coexisting deterministic and stochastic optima if *µ*_1_ < *µ* < *µ*_2_, and a single stochastic optimum if *µ* > *µ*_2_. Thus, making protein more costly is associated with lower expression levels as well as the transition toward a stochastic regime of expression. In the leaky case, the aforementioned holds with one caveat: it is optimal not to express protein if *µ* > *µ*_4_. Thus, if the cost of expressing the protein is too high, it is optimal to switch it off despite the protection it confers.

Deterministic and stochastic optima present contrasting strategies to use the protein to survive treatment. In the former, the protein is maintained at high levels throughout the population, so as achieve a high probability of survival in all cells. In the latter, the population expresses the protein in random bursts, with a small subpopulation of cells being sufficiently protected at any given time. The bifurcation diagrams reveal hysteresis in transitions between stochastic and deterministic optima. Consequently, we observe an all-or-none property in gene expression noise: optima are either deterministic or highly stochastic; indeed, stochastic optima are characterised by large burst sizes and low mean values, leading to large coefficients of variation CV^2^ = *b/x*. Our results thus suggest the existence of a noise floor for resistance driven by stochastic gene expression.

## Appendix A Asymptotics of the cost function

### The limit of *b* → 0 with *x* fixed (linear noise approximation)

Taylor expanding the survival probability around the protein mean gives:

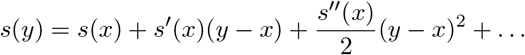

Taking the expectation with respect to the protein distribution gives

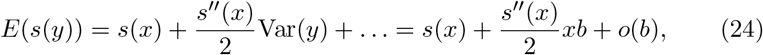

in which we substituted the variance of the gamma distribution [44]; higher-order central moments of the gamma distribution are *o*(*b*). Taking the logarithm of (24) yields

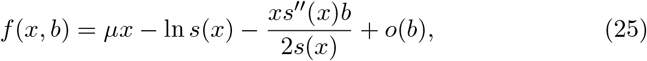

in which the *O*(1) term is equal to the cost of a deterministic expression (13) and the *O*(*b*) term gives the cost derivative (15) due to a stochastic perturbation of the deterministic expression.

#### The limit of *x* → 0 with *b* fixed

This is equivalent to taking *a* = *x/b* → 0.

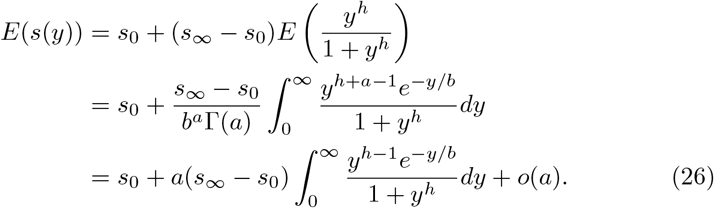

Taking the logarithm of (26) leads to

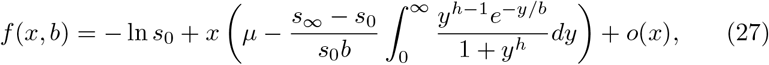

in which the *O*(1) term is equal to the cost of the trivial state (22) and the *O*(*x*) term gives the cost derivative (23) due to a stochastic perturbation of the trivial state. The derivative as function of mean burst size *b* can change sign depending on the burden rate *µ* (Figure 5).

**Figure 5:**
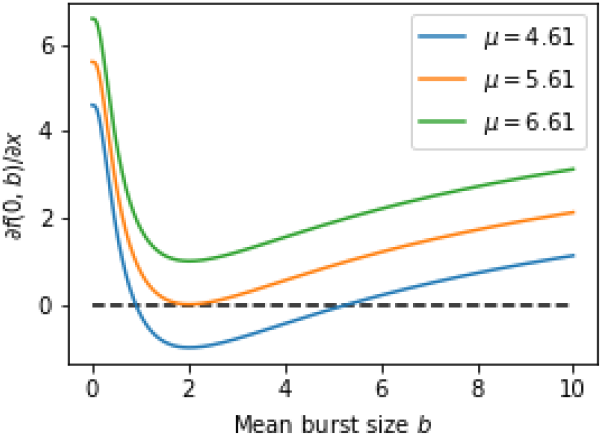
The cost derivative *∂f* (0, *b*)*/∂x* has zero, one, or two zeros depending on the burden rate *µ*. Parameter values are *h* = 4, *s*_0_ = 0.05, *s*_*∞*_ = 1.

## Appendix B Parameter continuation

We provide a specialised algorithm for the *µ*-continuation of stochastic critical points of the cost function (10). It is easier to implement and is expected to be more efficient than general approaches [45, 46].

Inserting (6) into (10) and simplifying give

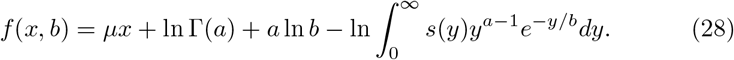

The burst frequency *a* in (28) is understood to be a function of the protein mean *x* and the burst size *b*:

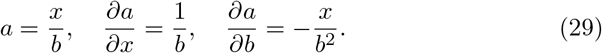

Differentiating (28) gives

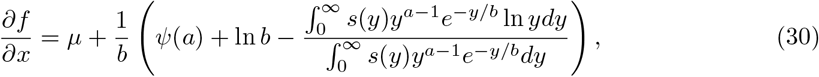

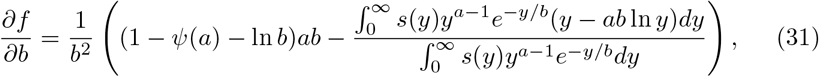

where *ψ*(*a*) = *d* ln Γ(*a*)*/da* is the digamma function. The bifurcation parameter *µ* features only as an additive term in (30) and is absent from (31). This simplicity facilitates an easy calculation of a bifurcation diagram as described below.

The critical point equations (17) are solved in the extended domain 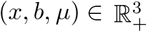 (gene expression and bifurcation parameters) in the following steps:

1. Pick a value *b* > 0.
2. Solve numerically in *a* = *a*(*b*) the critical equation *∂f/∂b* = 0, i.e.

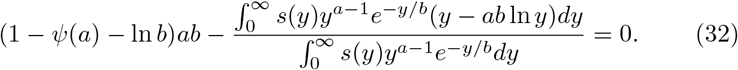
3. Set *x* = *x*(*b*) = *a*(*b*)*b*.
4. Use the second critical equation *∂f/∂x* = 0 to calculate *µ* = *µ*(*b*) for the given pair of *a* = *a*(*b*) and *b*, i.e. set

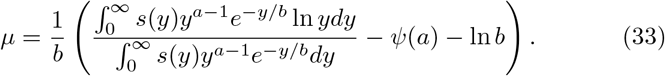
5. Perturb the value *b* → *b* + *ε* and go to Step 2.

Equation (32) is solved practically using the Newton method, which requires an initial guess for the solution *a* = *a*(*b*). In the first iteration, we assume that such a guess is available (e.g. from a visual inspection of the cost function). In the second iteration, we use the solution *a*(*b*) from the first iteration as the initial guess for the new solution *a*(*b* + *ε*), which itself becomes the initial guess in the next iteration, etc.

The above procedure constructs a curve (*x, b, µ*) = (*x*(*b*), *b, µ*(*b*)) parametrised by *b* > 0 in the extended domain of the cost function; (*x*(*b*), *b*) is thereby a critical point of the cost function for *µ* = *µ*(*b*). Plotting *x*(*b*) and *b* against *µ*(*b*) for a range of *b* values visualises how stochastic critical points (*x, b*) depend on the bifurcation parameter *µ*; such plots are shown in the main text.

